# Search behavior of individual foragers involves neurotransmitter systems characteristic for social scouting

**DOI:** 10.1101/2020.12.30.424710

**Authors:** Arumoy Chatterjee, Deepika Bais, Axel Brockmann, Divya Ramesh

**Author notes:** **Correspondence**: Divya Ramesh.

## Abstract

In honey bees search behavior occurs as social and solitary behavior. In the context of foraging, searching for food sources is performed by behavioral specialized foragers, the scouts. When the scouts have found a new food source, they recruit other foragers (recruits). These recruits never search for a new food source on their own. However, when the food source is experimentally removed, they start searching for that food source. Our study provides a detailed description of this solitary search behavior and the variation of this behavior among individual foragers. Furthermore, mass spectrometric measurement showed that the initiation and performance of this solitary search behavior is associated with changes in glutamate, GABA, histamine, aspartate and the catecholaminergic system in the optic lobes and central brain area. These findings strikingly correspond with the results of an earlier study that showed that scouts and recruits differ in the expression of glutamate and GABA receptors. Together, the results of both studies provide first clear support for the hypothesis that behavioral specialization in honey bees is based on adjusting modulatory systems involved in solitary behavior to increase the probability or frequency of that behavior.

## 1 Introduction

In honey bees, searching for food sources and collecting the food are performed by two different worker groups, scouts, and recruits (Lindauer, 1952; Seeley, 1983; zu Oettingen-Spielberg, 1949). Depending on the season and colony state, 5 to 25% of the foragers are scouts and all the others are recruits. Scouts search for new food sources every day; and recruits continue to visit a known food source for as long as the food source provides sufficient good quality food (Seeley, 1983, 1995). Based on these behavioral differences, it was proposed that scouts are similar to novelty seekers in birds and humans (Liang et. al., 2012, 2014).

In contrast, recruits only search for a food source at the beginning of their foraging career or when they decide to switch a food source, which does not occur very often during their short life (von Frisch, 1965; Seeley, 1983, 1995). After following a dance, the recruits search for the location of the food using path integration information indicated by the dance that they had followed (Riley et. al., 2005; von Frisch, 1965). Reaching the vicinity of the area indicated by the dances, they start searching for the food sources likely using odor cues perceived on the dancer (Farina et. al., 2005; von Frisch, 1965) as well as visual and floral scent cues of flowers in the area (Bell, 1990; Rachersberger et. al., 2019). Apart from that, recruits, i.e., foragers continuously foraging at a food source, have been shown to initiate a search behavior when the training feeder was experimentally removed (Reynolds et. al., 2007; Chatterjee et. al., 2019, Srinivasan et. al., 1997; Townsend-Mehler et. al., 2011; Townsend-Mehler and Dyer, 2012). The search behavior consists of increasing loops centering around the location where they expected the feeder with increasing radius and an orientation in the hive-feeder axis before they return to the hive (Reynolds et. al., 2007). Furthermore, similar experiments with an unscented feeder in a flight tunnel suggest that honey bee foragers predominantly use path integration and landmark memory when searching for a missing feeder (Srinivasan et. al., 1997).

In this study we explored two phenomena. Reynolds et. al. (2007) only observed the trajectory of flights when the bees experienced the missing feeder for the first time. Thus, the question remained whether the bees continue to make additional search flights, and if so, whether foragers show differences in their search behavior. We measured flight and hive durations of individually identified foragers for about 2 hours after removing the feeder. The temporal data were sufficient to describe changes in the behavior over time as well as distinguish individual differences in behavior. In addition, we were interested to know whether this search behavior might be regulated by neuromodulator systems involved in scouting behavior linking individual behavior to social division of labor and behavioral specialization (Liang et. al., 2012). There is growing evidence that behavioral specialists might be temporarily tuned in to a specific brain and behavioral state that occurs in any individual of the species when they perform the corresponding solitary behavior (Toth et. al., 2005; Alaux et. al., 2009; Shpigler et. al., 2017). Comparing brain gene expression in scouts and recruits and manipulative experiments Liang et. al. (2012) identified that changes in catecholamine (*DopR1*), glutamate (*Eaat-2*, *Vglut*, *Glu-RI*), and γ-aminobutyric acid signaling (*Gat-a*) are associated with scouting behavior. Furthermore, manipulative experiments confirmed that glutamate and octopamine treatment increased, and dopamine antagonist treatment decreased the likelihood of scouting (Liang et. al., 2014, 2012). Thus, we were specifically interested whether these neurotransmitter systems are also involved in search behaviors performed by regular foragers, i.e., recruits, when they do not find a known feeder. We used mass spectrometry measurements (Ramesh and Brockmann, 2019) to test whether the search behavior of recruits which was induced by the removal of a visited feeder is associated with short-term changes in neuromodulators involved in social scouting. The titer measurements were done for two brain areas of behaviorally characterized individual foragers: the central brain comprising the central complex and the mushroom bodies, which have been demonstrated to be involved in visual navigation including path integration and landmark learning (Kamhi et. al., 2020; Buehlmann et. al., 2020; Stone et. al., 2017, Seelig and Jayaraman, 2015) and the optic lobes pre-processing the visual information used for navigation and landmark memory (Brockmann and Robinson, 2007; Yilmaz et. al., 2019; Zeller et. al., 2015).

## 2 Results

### 2.1 Absence of an expected feeder elicited a series of search flights and subsequent cessation of foraging

Honey bee foragers (BE 1: n=16, 2015 and n=16, 2020) that had continuously visited a feeder for a few hours immediately initiated a search when they did not find the feeder at the expected location (Figures 1A-C). Already the mean duration of the foraging trip when they did not find the feeder (FS, combined foraging/search flight; Table S1) was significantly longer than the mean duration of the foraging trip (FS, Figures 1D, E). In contrast, the mean duration of the hive stay after this first unsuccessful trip (HFS) was as short as those after the previous regular foraging trips (HF; Figures 1F, G). One of the foragers directly stopped foraging after the FS flight (BeeID: E26; Figure 1C), whereas all the other foragers performed one to four additional search flights (Figures 1B, C). All foragers stopped foraging within 100 mins after the removal of the feeder. The mean duration of the consecutive search flights was relatively consistent (S: 12.37 ± 6.02 min) and lasted about 3 times longer than the mean duration of the foraging trips (F: 3.51 ± 1.02 min; Figures 1D, E; GLMM gamma family and generalized linear hypothesis test; see also supplementary data file S1 for details of the GLMM and GLHT results). In contrast to the search flights, the mean duration of the intermittent hive stays (HS1 - HS3) increased with the number of search trips (Figures 1F, G; GLMM gamma family and generalized linear hypothesis test; see also supplementary data file S1 for details of the GLMM and GLHT results).

**Figure 1.**
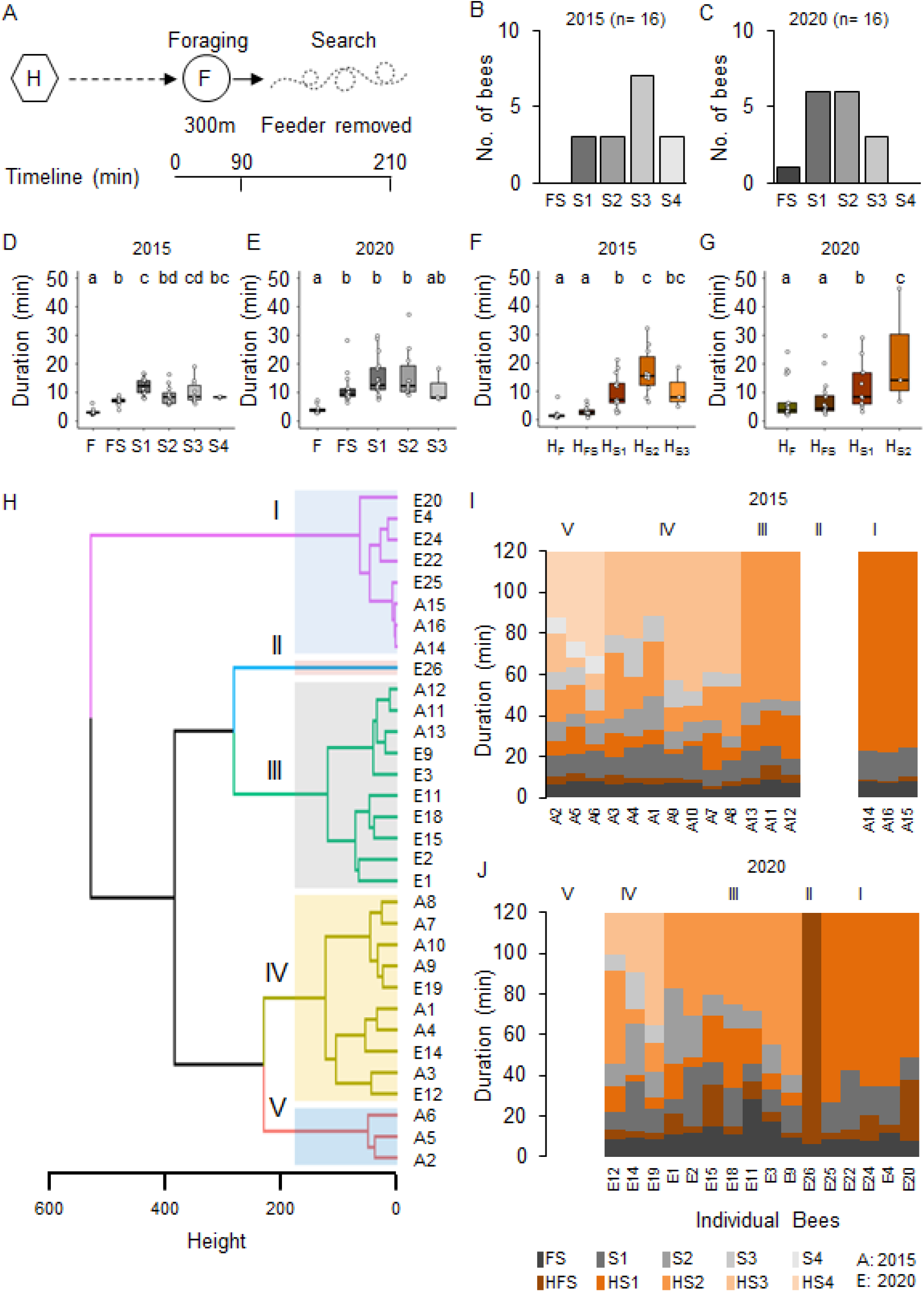
Removal of the feeder led foragers to perform search flights. (A) Experimental design to study the dynamics and persistence of search behavior and after foragers were confronted with the absence of a known feeder. Individually marked foragers are allowed to visit the feeder at 300m distance from the hive for an initial 1.5 hrs. The feeder is removed, and the outbound flight activity of the marked foragers is monitored at the hive entrance for the following 2 hrs. (B-C) Search flights following feeder removal for two colonies. (D-E) Increase in hive-to-hive duration and (F-G) duration of hive stays before and after feeder removal. (H) Hierarchical clustering of foragers based on search behavior sequence. The maximum average silhouette width 0.58 gave a five-cluster solution with agglomerative coefficient 0.93. (I-J) Search behavior sequences for foragers along with cluster information for two colonies.

In an additional control experiment (BE 2) in which we put the feeder back after 1 hour, foragers landed on the feeder as soon as it was opened (Figure S1). This finding suggests that the search flights were more or less restricted to the close vicinity of the expected feeder location and the foragers were not searching for any other food location.

### 2.2 Individual foragers showed different search phenotypes

Cluster analysis based on the number and temporal dynamics of the search flights and hive stays identified five different search phenotypes independent of the behavioral experiment (I-V; Figures 1H-J; see Figure S2A for optimum number of clusters). Cluster 1 includes bees that stopped foraging after the first search flight (S1; n=3, 2015; n=5, 2020) and cluster 2 includes the single forager that already stopped foraging after FS (BeeID: E26, 2020). Foragers in Cluster III (n=3, 2015 and n=7, 2020) made 2 search flights and Cluster IV (n=7, 2015 and n=3, 2020) made 3 search flights within the observation period. Cluster III and IV formed the largest groups each with 10 bees. Cluster V comprised 3 foragers (all in 2015) which performed 4 search trips (see supplementary data file S1 for details). As the number of search flights is the parameter with the strongest impact, the different clusters present behavioral phenotypes that vary in their motivation to search and their persistence to continue foraging.

### 2.3 Search behavior led to a robust reduction of glutamate and GABA titers in the central brain

Neurotransmitter analysis of the brain parts (Figure S3) from different foragers (Table S2) was done in multiple batches, each containing samples of bees from all behavioral groups (see supplementary data file S1). The batch identity was added as a random factor in the statistical model. Comparing neurotransmitter titers in the central brain (CB) between foragers caught during foraging, searching for the feeder, or revisiting the feeder (Figure 2A), we found robust differences for glutamate and GABA (Figures 2B, C). Foragers that had experienced the absence of the feeder for the first time (FS) and were caught as they were leaving for their first search trip already showed significantly lower GABA titers in the CB (Figure 2C; decrease by 29.7 +/− 10.3 ng, p=0.025) than successful foragers. In contrast, glutamate titers in the CB declined after a first search flight (Figure 2B; decrease by 242.2 +/− 80.8 ng, p=0.018). Further, glutamate levels continued to linearly decrease with the number of search trips (Figure 2D; decrease by 111.7 +/− 34.8 ng with every search flight, p=0.001). Similarly, GABA levels also showed a significant linear decrease, however, the largest reduction occurred during the FS trip and the following hive stay (Figure 2E; decrease after first experience by 29.5 +/− 10.6 ng, p=0.032).

**Figure 2.**
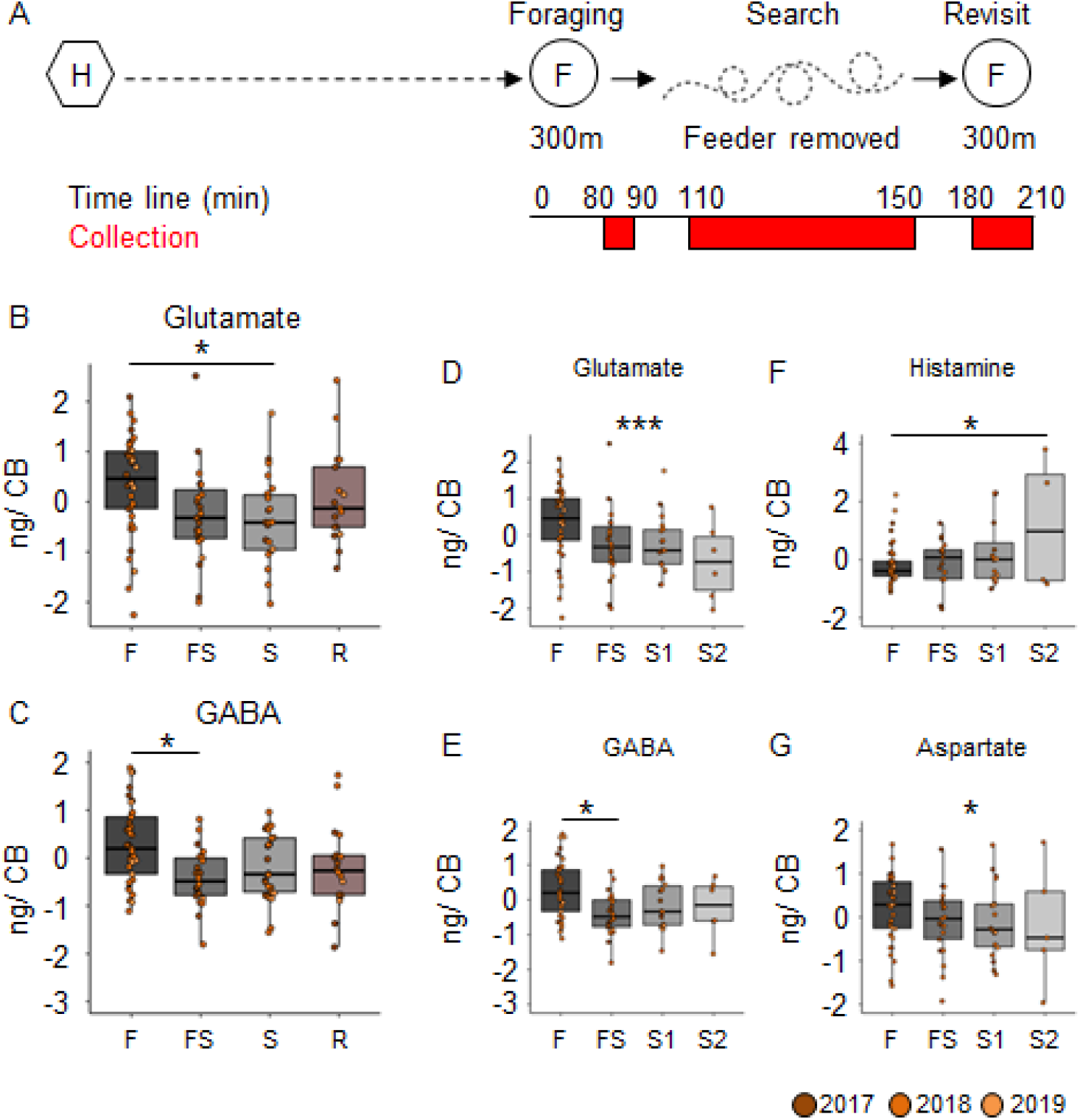
Search behavior is correlated with reduced GABA and glutamate titers in the central brain. (A) Experimental design to collect foragers for mass spectrometric analysis of brain neurotransmitter titers. Individually marked foragers are allowed to visit the feeder at 300m distance from the hive for an initial 1.5 hrs (foraging phase). The feeder is removed for 1 hr (search phase) and reinstalled for 1 hr (revisit phase). Foragers were captured at hive entrance during each behavior phase (marked red on experiment timeline). (B-C) Glutamate and GABA levels decrease in the central brain (CB) after bees experience a loss of their expected feeder. (D-E) A detailed look at the dynamics of change indicates that glutamate levels gradually but linearly decrease over increasing search trips (decrease by 112 ng per search flight, p-value = 0.001), but GABA levels only decrease post the first experience and stay that way. (F) Histamine (G) Aspartate (decrease by 87 ng per search flight, p-value = 0.031). The neurotransmitter values are scaled by the MS batch. *p<0.05; **p<0.01; ***p<0.001.

In addition to the changes in GABA and glutamate titers, we also found differences in the histamine and aspartate levels in the central brain samples (Figures 2F, G). Foragers with 2 search flights had significantly higher histamine levels than those that were foraging, and the histamine levels showed a significant linear increase with number of search flights (0.21 +/− 0.096 ng per search flight, p=0.025). In contrast, aspartate levels showed a significant linear decrease with number of search flights (decrease by 86.8 +/− 40.28 ng per search flight, p=0.034).

### 2.4 Restarting foraging led to an increase of glutamate and GABA titers in the optic lobes

In contrast to the CB, we did not detect any changes in neuromodulator titers in the optic lobes (OL) samples during the search flights (Figures 3A-D). However, when we reinstalled the feeder, the foragers, that had restarted foraging, showed significantly higher levels of glutamate and GABA than any other behavioral group (Figures 3A-D). Furthermore, we also found changes in the titers of other neurotransmitters and their precursors in the optic lobes after the bees restarted foraging (Figures 3E-O). Tyrosine and L-DOPA, but not dopamine, were significantly higher in foragers that had restarted foraging compared to the foragers visiting before the feeder was removed (Figures 3F, G). Tryptophan, aspartate, histamine, and serine were higher in restarted foraging compared to those that had performed search flights and those that foraged before the removal of the feeder (Figures 3K-O; supplementary data file S1). All neuromodulators, for which we detected a change, showed a significant increase in their titers due to revisiting the feeder. These dramatic changes were independent of the number of search flights (Figure S4).

**Figure 3.**
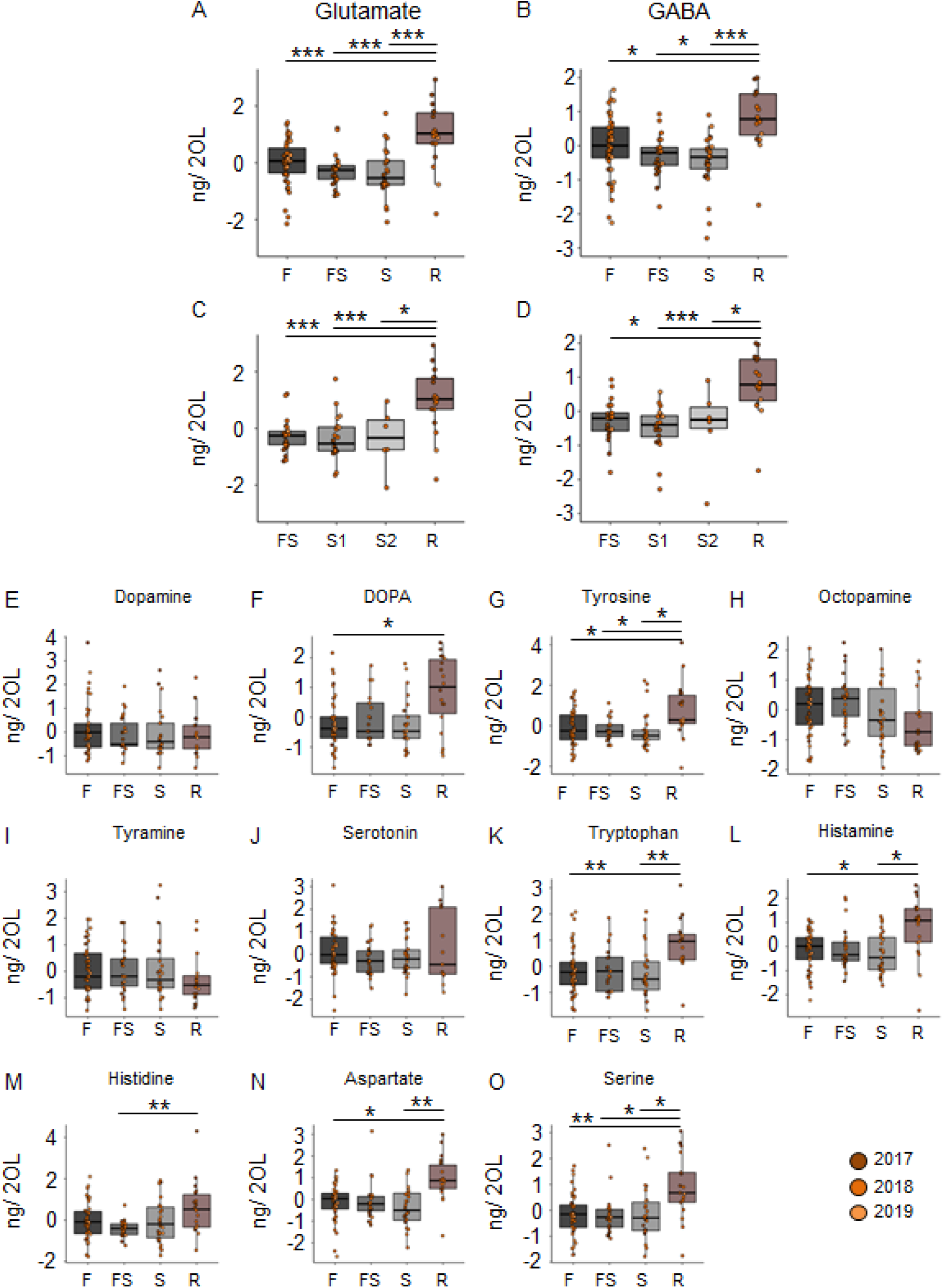
Reinitiation of foraging increases levels of Glutamate and GABA in the optic lobes. (A-B) Glutamate and GABA levels increase after bees start revisiting the feeder. There are no changes due to the experience of feeder loss. (C-D) A detailed look at the dynamics show that the number of search trips do not affect the modulator levels, but only the experience of the feeder does. (E-O) Replacement of the feeder causes abrupt and global changes in multiple modulators in the OL. The neurotransmitter values are scaled by the MS batch. *p<0.05; **p<0.01; ***p<0.001.

### 2.5 Forager search phenotypes show differences in the titers of HA, Octopamine and L-Dopa

Based on our findings that foragers differ in their motivation to search and their persistence to forage, we performed a cluster analysis on the individual temporal search dynamics of the collected foragers. Of course, the behavioral data of the collected foragers do not allow a clear identification of the search phenotype for all collected foragers because we collected them during their search behavior instead of after they had stopped leaving the hive. However, we were able to identify a group of foragers that performed several search flights with short intermittent hive stays (Cluster IIIe, Figures 4A, B) and foragers that already showed a long hive stay after the foraging/search flight (FS, Cluster I, Figures 4A, B) before they were collected. With respect to our behavioral analyses, these two search phenotypes strongly differ in their motivation to search. Only for the collection experiment performed in 2018, we found a sufficient number of foragers with different search phenotypes for a comparison of the neurotransmitter levels (Figures 4A, B).

**Figure 4.**
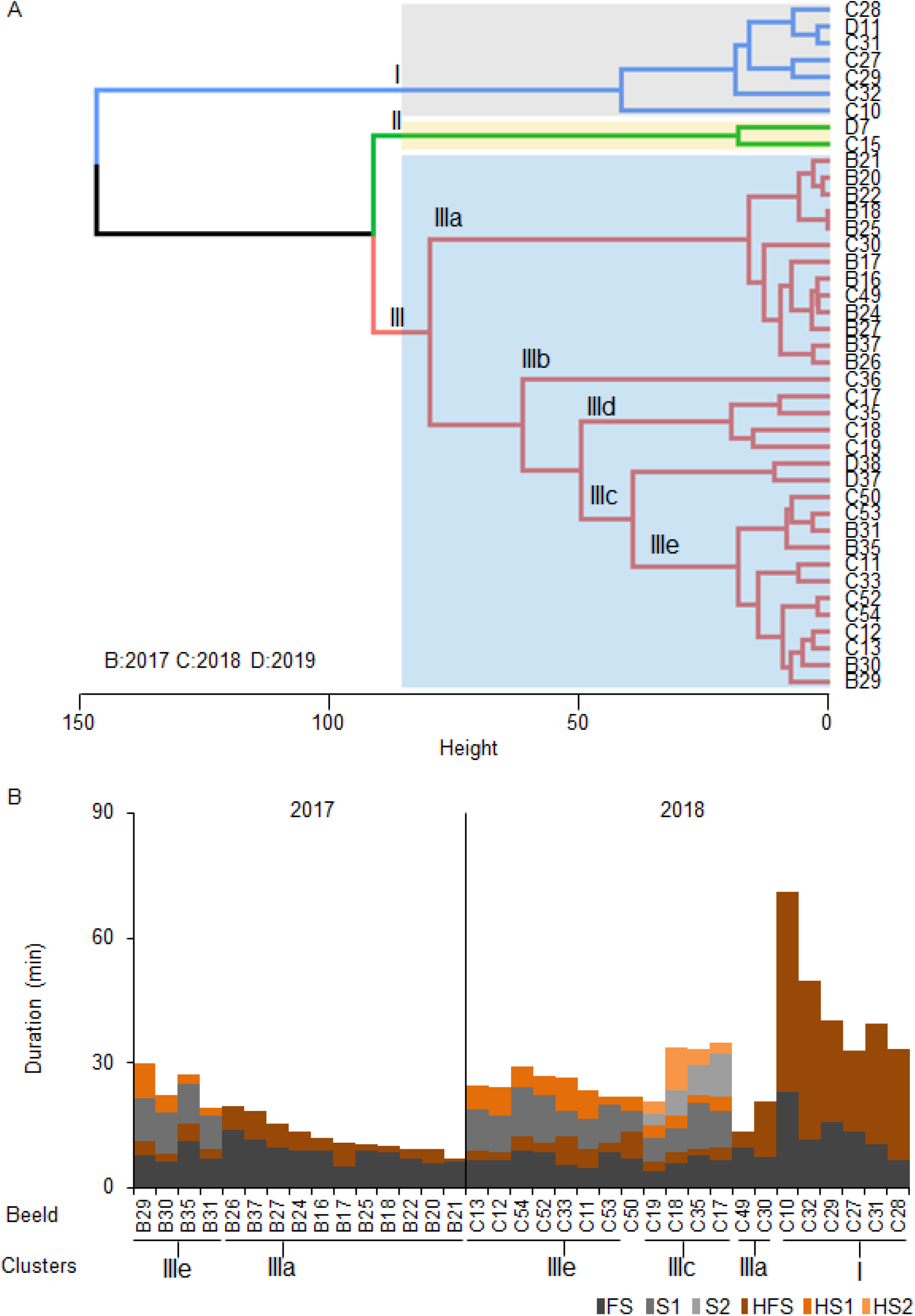
Phenotypes of the collected foragers. (A) Hierarchical clustering of foragers collected during search phase based on search behavior sequences. The maximum average silhouette width 0.52 gave a three-cluster solution with agglomerative coefficient 0.94. (B) Search behavior sequences for the foragers from the collection experiments along with cluster information. Only bees used for cluster analysis of neurotransmitter titers are shown.

Foragers of the cluster IIIc that had performed 2 search flights with short intermittent hive stays showed significantly higher DOPA and HA levels and significantly lower octopamine levels in the central brain samples compared to one or more groups of foragers with fewer search flights (Figures 5A-C). Foragers of the cluster IIIc also showed a lower level of aspartate in the optic lobes as compared to foragers of the other three clusters (Figure 5E). This difference was also observed between similar phenotypes in the 2017 collection (Figure 5D).

**Figure 5.**
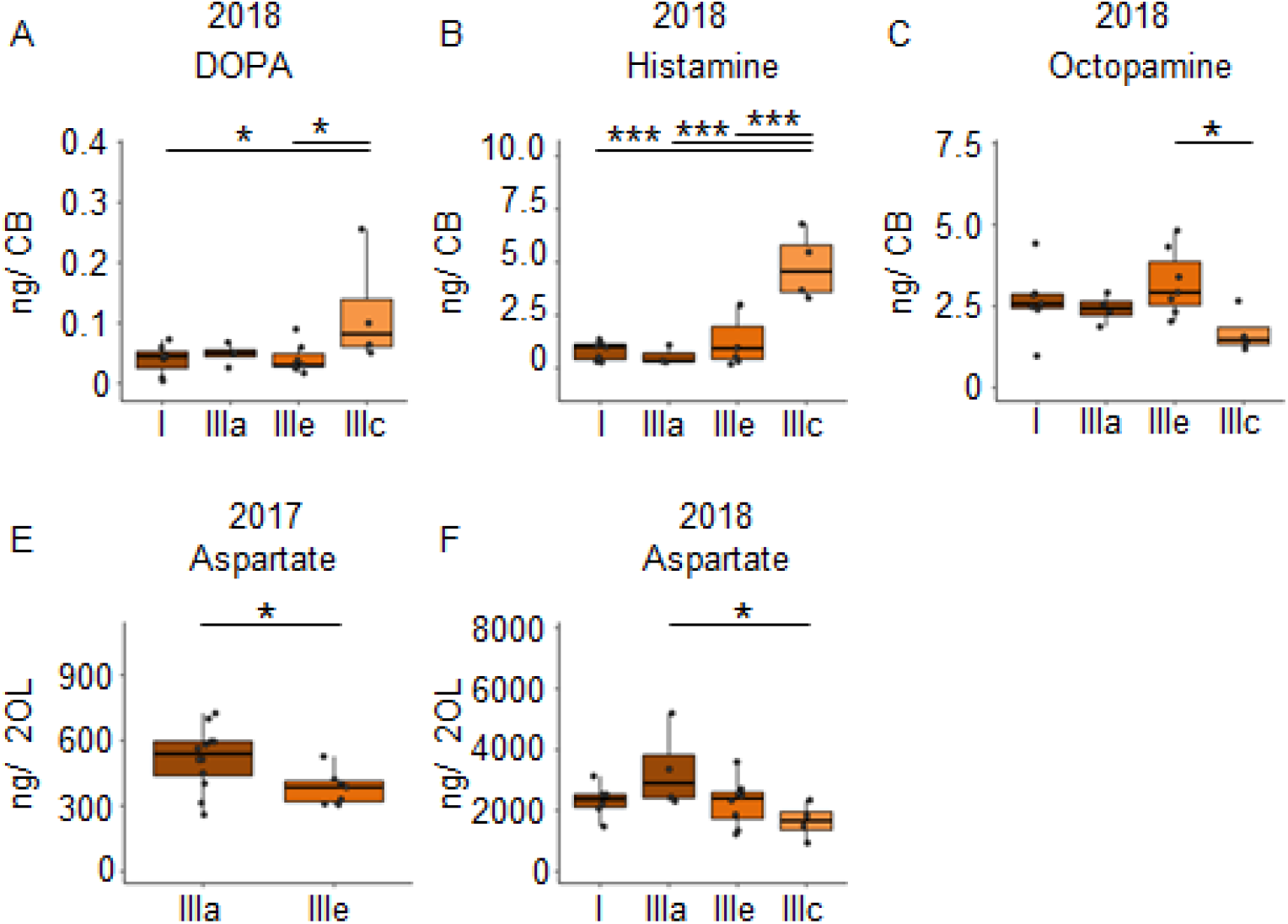
Search intensity negatively correlates with OA and positively correlates with DOPA and HA titers in the CB. (A-B) DOPA and Histamine levels show an increase with increased search trips. (C) Octopamine levels decrease with increased search trips. (D-E) Foragers most motivated to continue search flights showed decreased levels of aspartate in the optic lobes. In addition, bees that performed the FS trip but are phenotypically different in their hive stay times show differences in aspartate levels, though non-significant. Bees without detailed behavior data were added to the phenotype analysis by comparing available relevant behaviors. *p<0.05; **p<0.01; ***p<0.001.

### 2.6 Colonies vary significantly in their neurochemical signatures

Neurochemical content from the CB and OL were quantified from foragers from three different colonies and used for analyzing differences in behavior. In addition to finding changes related to search and restarting of foraging, we found significant differences across colonies as well. A PCA analysis of the CB and OL titers showed that the colonies clustered separately (Figures 6A, B), and that more than 50% of the variance in transmitter content is explained by the colony differences alone. Individually as well, transmitters showed significant differences between colonies (Figures 6C-O and Figure S5). Specifically, the colony used for CE 1 had lower amounts of transmitters in general, in comparison to the other colonies. In the OL, 12 out of 14 transmitters in CE 1 were significantly lower than CEs 2 and 3, while in the CB, 10 transmitters were significantly lower than at least one of the other colonies (Figure S5). In spite of the large differences in transmitter titers due to the colony identity, we were still able to detect the changes in neurochemicals due to the behavioral state.

**Figure 6.**
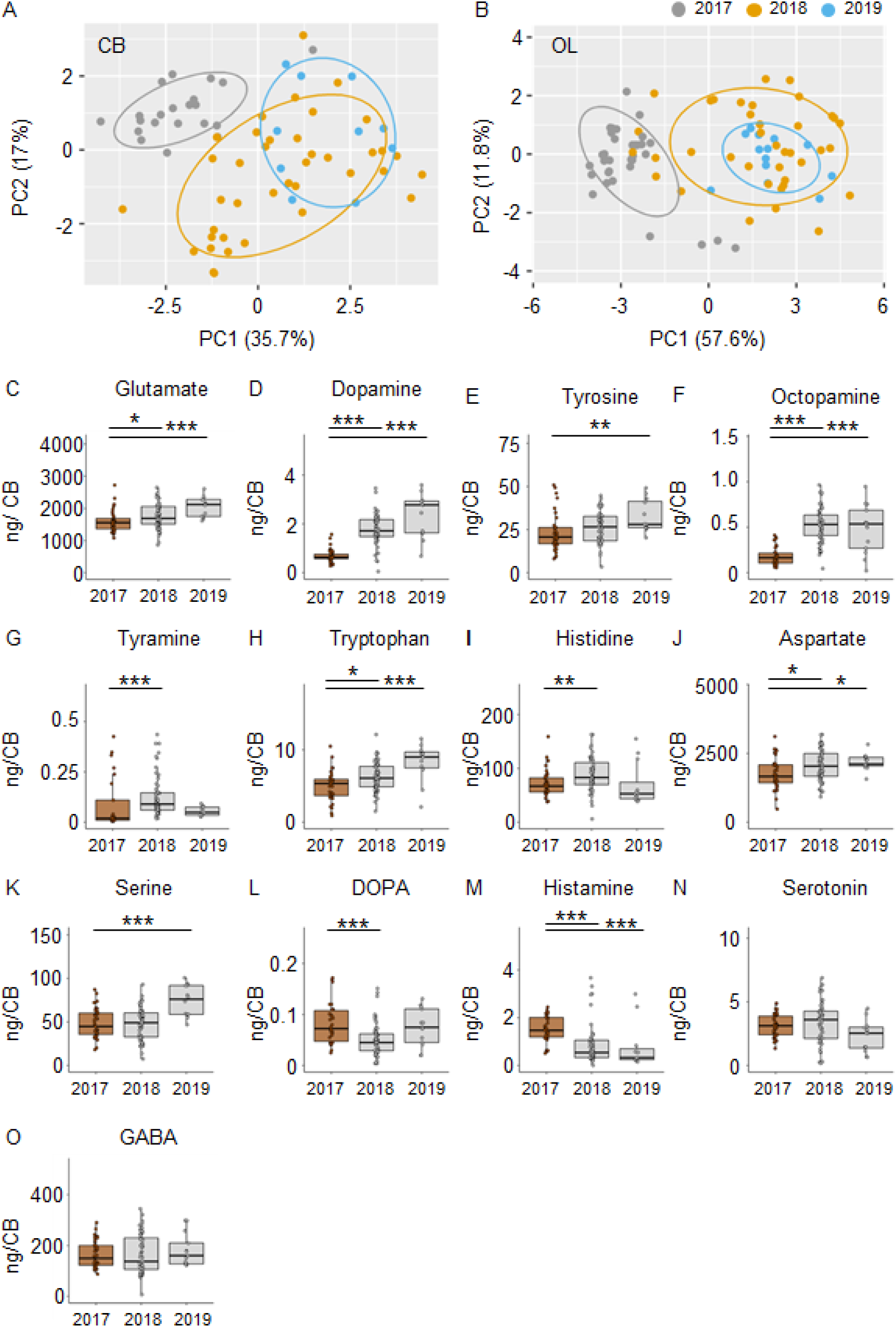
Neuromodulators vary significantly between different colonies. PCA of modulator titers in the CB (A) and OL (B) show >50% variance explained by colony membership. (C-O) Individual modulators vary significantly between the different colonies. In general, the 2017 colony shows a lower amount of most transmitters than the 2018 and 2019 colonies. Only the differences between the 2017 and the other two years are shown. *p<0.05; **p<0.01; ***p<0.001.

## 3 Discussion

The principal result of our study is that glutamate and GABA titers in the central brain region (comprising mushroom bodies, central complex and adjacent protocerebral areas) decreased during continuous search behavior for a previously visited but absent feeder. This finding corresponds with the fact that the brains of scouts show a higher expression of genes involved in glutamate and GABA signaling. This correlation suggests that the behavioral specialization is based on genomic mechanisms that modulate signaling mechanisms used in regular search behavior.

### 3.1 Search behavior of foragers that do not find a known feeder at the expected location

Our analyses of the temporal dynamics of flight and hive stay durations of foragers that did not find a known feeder at the expected location suggest that the initiated search behavior consists of a few to several search flights of relatively similar duration (BE 1: range 1-4 search flights; mean 12.37 ± 6.02 min; range = 5.79 - 37.3 min, Figures 1D,E), and intermittent hive stays that get longer with the increasing number of search trips (BE 1: 13.43 ± 9.37 min, range= 2.33 - 46.4 min; Figures 1F,G). The duration of the hive stay appears to be a good predictor of the probability to stop the search (and foraging) for a period of time. In the additional control experiment (BE 2), in which we reinstalled the feeder after 1 hour, foragers almost immediately started landing on the feeder, indicating that the bees’ search flights were more or less restricted to the vicinity of the feeder location.

These observations nicely correspond to findings of other studies in which similar experiments were performed. Radar tracking experiments showed that foragers that did not find the expected feeder started flying in loops around the expected location of the feeder and after some time returned to the hive (mean duration 4.49 ± 2.44 min, Reynolds et. al., 2007). Furthermore, the search flights were mostly oriented in the hive to feeder direction (Reynolds et. al., 2007). Al Toufailia et. al., (2013), reported that foragers trained for a few days revisited an empty feeder at an average of 4.29 ± 4.47 trips (range: 0–25) over a 6-hour recording period. The persistence to revisit the temporarily unrewarded feeder, measured as number of trips and total duration of trips, correlated with previous foraging experiences, e.g., duration of feeder availability and profitability, as well as season, which affects colony food stores (Al Toufailia et. al., 2013; Townsend-Mehler et. al., 2011; Townsend-Mehler and Dyer, 2012). Furthermore, trained foragers were found to continue to visit an emptied feeder for a few days. (1.89 ± 1.56 days, range 0–7 days, Al Toufailia et. al., 2013; see Table S3).

Together all these studies indicate that the search behavior elicited by the absence of a known feeder induces a search behavior for this feeder. These foragers are not searching for a new food source or food location as scouts do. Only after repeated unsuccessful visits over a few days, is it reported that the majority of foragers might start searching for a new food source, and that too, most likely only after following dances (Biesmeijer and de Vries, 2001; Seeley, 1983). None of the searching bees were found to follow any unmarked dancer during their hive stay within the observation period.

### 3.2 Individual variation and search phenotypes

In addition to the description of the general temporal dynamics of this search behavior, cluster analysis showed that the individual foragers visiting the same feeder varied in the intensity of search behavior or persistence in revisiting the feeder location. The strongest search response is characterized by fast repetition of search flights which includes short intermittent hive stays. The weakest response was characterized by 1-2 search flights with long intermittent hive stays. These differences are likely based on variations in the behavioral state depending on previous experience and genotype.

In an earlier study, our lab reported that there are consistent long-term individual differences in the dance activity among foragers visiting the same feeder. Interestingly, these differences were, at least to some degree, dependent on the composition of the group (George et. al., 2020). Furthermore, the individual variation in dance activity correlated with expression differences in the foraging gene (*Amfor*). Similarly, linear discriminant analysis of the brain gene expression pattern of individual scouts and recruits showed a separate but overlapping distribution, suggesting a more quantitative than qualitative difference between these phenotypes (Biesmeijer and de Vries, 2001; Liang et. al., 2012). For the future, it will be interesting to explore whether differences in foraging, dance and search activity among foragers visiting the same feeder are correlated and based on the same physiological processes or not. In addition, it would be interesting to see whether scouts resemble one of these forager phenotypes or represent a separate one.

### 3.3 GABA and glutamatergic systems are involved both in search behavior of recruits and scouts

The comparison of foragers with different numbers of search flights suggest that glutamate and GABA titers continuously decrease with the number of search trips in the central brain (i.e., mushroom bodies, central complex and surrounding protocerebral brain areas). In contrast, glutamate and GABA titers in the optic lobes did not change during search but showed an abrupt increase when the foragers had started revisiting the feeder. This kind of rebound increase in titers was not observed for the central brain region; moreover, the titers were still lower compared to the foraging group at the beginning of the experiment.

Liang et. al., (2012) reported a higher expression of several glutamate and GABA receptor and transporter genes in the brains of scouts compared to recruits. In addition, treatment experiments with monosodium glutamate (MSG) increased scouting behavior. Although we do not know the exact function of glutamate and GABA in search behavior (Carr-Markell and Robinson, 2014; Cook et. al., 2019; Filla and Menzel, 2015; Locatelli et. al., 2005), the comparison of scouts and recruits and our studies on search behavior in foragers (i.e., recruits) strongly suggest that these neuromodulators have an important function in search behavior in general. Changes in the glutamate and GABA signaling appear to be major physiological underpinnings of the behavioral specialization of scout bees. Furthermore, this molecular mechanism might not be unique to honey bees, as it was found that glutamate receptors are also upregulated in scouts of *Temnothorax* ants (Alleman et. al., 2019).

Finally, one of the most original recent molecular studies in honey bees showed an increase in activity in GABA-ergic neurons of the optic lobes during re-orientation flights in which the foragers learn the hive entrance and hive location (Degen et. al., 2018; Kiya and Kubo, 2010). In our behavioral experiments the foragers that found the feeder again also performed learning flights involving circling over the feeder (Figure S6; Lehrer, 1991). Thus, the abrupt increase in GABA titers in the optic lobes in the revisiting foragers might be related to the phenomenon described by Kiya and Kubo (2010). GABA-ergic neurons in the optic lobes of *Drosophila*, for example, are involved in tuning the sensitivity and selectivity of different visual channels (Keleş et. al., 2020; Keleş and Frye, 2017). Similarly, glutamate signaling in the optic lobes is involved in shaping object recognition and directional motion vision (Bicker et. al., 1988; Rossi et. al., 2020; Sinakevitch and Strausfeld, 2004).

In addition to the differences in the glutamate and GABA titers, we found changes in histamine and aspartate. Histamine levels in the CB increased with the intensity of search (Figures 2F and 5B), and in the OL, they increased due to the re-initiation of foraging (Figure 3L). Aspartate was found to decrease linearly with increasing search trips in the central brain (Figure 2G) as well as in the optic lobes (Figures 5D, E). Later, during the re-initiation of foraging, aspartate levels in the OL increased (Figure 3N). There is growing evidence that neuromodulation in the optic lobes plays a significant role in selecting and adjusting visual processing according to the behavioral context (Cheng and Frye, 2020). Our results suggest that HA and aspartate, which showed significantly higher titers in the optic lobes of revisiting foragers, might play an important role in modulating visual processing (Hamanaka et. al., 2012; Sinakevitch and Strausfeld, 2004; Thamm et. al., 2017; Ziegler et. al., 2013). The changes in the central brain are more difficult to interpret. Previously, we reported that aspartate levels increased globally during anticipation of food (Ramesh and Brockmann, 2019). It is therefore likely that aspartate and histamine play a role in regulating foraging and motivation (see Torrealba et. al., 2012), although this remains to be investigated.

### 3.4 Forager search phenotypes show differences in the titers of HA, Octopamine and L-Dopa

Comparison of search phenotypes revealed that foragers with a high intensity of search behavior (several successive search flights with short intermittent hive stays, Cluster IIIe) had higher DOPA and HA levels and lower OA levels in the central brain region than foragers with less intense search behavior. These differences in the levels of neurotransmitters among search phenotypes could be a result of a higher degree of neural signaling activity or differences in the baseline levels of neurotransmitter levels among search phenotypes.

Liang et. al., (2012) reported that octopamine treatment resulted in a weak but significant increase in scouting behavior and that scouts showed a higher expression of the *Octβ2R* receptor. Interestingly, they also found that a dopamine antagonist treatment inhibiting dopamine signaling caused a significant decrease in scouting, but their molecular data indicated that dopamine signaling might be downregulated in scouts. The two dopamine receptors, *AmDopR1* and *AmDopR2* showed a lower whole brain expression in scouts compared to non-scouts (Liang et. al., 2012). More recently, Linn et. al., (2020) showed that foragers treated with octopamine revisited an emptied feeder more often than a sham-treated control group. More importantly, they preferred the known but emptied feeder instead of searching for a new feeder indicated by other nestmates, suggesting that octopamine might increase foraging activity in the sense of persistence (or probability of leaving the hive) but not in a specific sense of searching (Barron et. al., 2002; Barron and Robinson, 2005; Wagener-Hulme et. al., 1999). Recently, Cook et. al., (2019) reported that the brains of scouts showed higher tyramine levels than those of recruits. However, similar to octopamine, the function of tyramine might not be directly involved in search behavior but foraging and flight activity, as suggested by QTL studies on honey bee foraging behavior (Hunt et. al., 2007).

### 3.5 Colonies differ in neuromodulator levels as well as in search phenotypes

In our study, we found that greater than 50% of the variance in neurotransmitter titers were due to the identity of the colony from which the foragers were caught (Figures 6A, B). An interesting observation was that the foragers of colony CE 1 showed lower titers compared to foragers of CEs 2 and 3 for almost all neuromodulators tested (Figures 6C-O and Figure S4). These were bees that were housed in an observation hive. Many previous studies on honey bees reported variation in brain neurotransmitter and neuromodulator titers with colony state and season (Božič and Woodring, 1998; Harris and Woodring, 1992). The colony hosted in the observation hive likely differed from the others in the density and crowding of bees, as well as in the available food stores (impacting the hunger state), both of which are known to affect the neurochemical composition of the brain (Hewlett et. al., 2018; Mayack and Naug, 2015). In spite of the large differences in transmitter titers due to the colony identity, we were still able to detect the changes in neurochemicals due to the behavioral state.

Colonies also differed in the composition of search phenotypes. In the behavior experiment (Figures 1I, J) as well as in the collection experiment (Figure 4B), performed over 5 years, we found that colonies differed in the presence and relative composition of phenotypes. These differences are likely due to colony conditions and forage availability modulating the foraging force.

### 3.6 Novelty seeking, glutamate, GABA and the honey bee brain

Liang et. al., (2012) suggested that the brain expression differences between scouts and recruits have something to do with novelty seeking as scouts are obviously searching for new food sites, and studies in vertebrates indicated that the identified neuromodulator systems (glutamate, GABA, and catecholamine) are involved in novelty seeking. Novelty seeking is certainly a complex behavior composed of different behavioral routines or modules, e.g., a specific flight pattern and increased visual and olfactory attention when searching for flowers. The differences in the changes in glutamate and GABA titers in the central brain region and optic lobes in our study might correspond to these different behavioral modules. Our finding that GABA titers increase with relearning the re-established feeder corresponds with the finding that GABA neurons in the optic lobes showed an increased activity during relearning the nest entrance and its surrounding after the hive had been experimentally relocated (Kiya and Kubo, 2010). Another question is whether glutamate and GABA initiate or modulate (enhance) search and scouting behavior (Palmer and Kristan Jr, 2011). A recent study in *Drosophila*, for example, showed that the majority of octopaminergic neurons are also glutamatergic and that both transmitters in these neurons affect the same and different behaviors, and thus might be involved in selection of behavioral modules (Sherer et. al., 2020). In honey bees, glutamate and GABA have mainly been studied in the context of learning and memory (Gauthier and Grünewald, 2012; Leboulle, 2012; Locatelli et. al., 2005; Shyu et. al., 2017; Xia et. al., 2005; Zwaka et. al., 2018). A study in ants aimed at identifying negative effects of increased monosodium glutamate consumption over several days showed a decrease in “precision of reaction”, a decrease in “the response to pheromones”, a decreased “impacted cognitive ability”, and “largely reduced learning and memory” (Cammaerts-Tricot and Cammaerts, 2016). Thus, the most plausible assumption at the moment might be that glutamate and GABA are involved in modulating brain circuits involved in search behavior and thus changing probabilities to perform behavioral routines involved in search behavior. The insect central brain region including mushroom bodies, central complex and adjacent protocerebral areas are involved initiating and selecting behaviors (Huber, 1955; Hulse et. al., 2020; Tsao et. al., 2018; Varga et. al., 2017), path integration and landmark learning (Buehlmann et. al., 2020; Kamhi et al., 2020; Seelig and Jayaraman 2015; Stone et. al., 2017); and the optic lobes in pre-processing the visual information used for navigation and landmark memory (Brockmann and Robinson, 2007; Yilmaz et. al., 2019; Zeller et. al., 2015).

For the future it would be interesting to identify neuron populations that are involved in search and scouting. One approach would be to perform double in-situ staining for neuronal activity-regulated genes and genes involved in glutamate and GABA signaling in brains of recruits and scouts caught during search behavior, as it was done for re-orienting foragers by (Kiya and Kubo, 2010; Sommerlandt et. al., 2019). Having identified neurons involved in searching as well as scouting, one could compare their expression patterns to identify the molecular changes underlying behavioral specialization at the cellular level. Regarding the neuronal mechanisms that determine scouts, it might be promising to first compare how scouts differ from recruits in the search behavior. For example, do scouts perform longer and more extended search flights? Subsequently, one could study whether the gene expression differences between scouts and recruits are based on changes in gene expression in the same cells or an extension of gene expression in different cells and identify in which brain areas these differences occur.

Honey bee foraging at an artificial feeder is one of the most fruitful and successful experimental assays in the study of animal behavior. All the fascinating behavioral and cognitive capabilities of honey bees have been identified using the feeder training assay (Giurfa, 2015; Seeley, 1995; von Frisch, 1965). As a behavior, foraging can be nicely dissected into different behavioral routines which can be studied separately, e.g., anticipating foraging in the hive (Ramesh and Brockmann, 2019; Reinhard et. al., 2004; Shah et. al., 2018), flying towards the feeder, food collection at the feeder (Brockmann et. al., 2009; Singh et. al., 2018), flying back to the hive, and recruiting nestmates with the dance (Barron et. al., 2007; Chatterjee et. al., 2019). In addition, there is increasing evidence that foragers vary in the intensity of each behavioral module (Cook et. al., 2019; George et. al., 2020; George and Brockmann, 2019; Jeanson and Weidenmüller, 2014). We suggest that detailed behavioral experiments combined with sophisticated molecular techniques will help to identify candidate neuronal mechanisms involved in elaborated behavioral and cognitive capabilities (Kiya and Kubo, 2011; Shah et. al., 2018; Sommerlandt et. al., 2019). Of course, decisive causal mechanistic studies will only be possible if genetically engineered honey bees are widely available (Değirmenci et. al., 2020; Kohno et. al., 2016; Roth et. al., 2019), or we might use *Drosophila* for comparative studies to identify neural underpinnings of some of the behavioral modules (Brockmann et. al., 2018; Kamhi et. al., 2017; Murata et. al., 2017; Reaume and Sokolowski, 2011).

## 4 Materials and Methods

### 4.1 Materials availability

*Apis mellifera* colonies were procured from HoneyDay Bee Farms Pvt. Ltd., Bangalore. All standards, formic acid (FA), hydrochloric acid (HCl), boric acid and ascorbic acid as well as reagents required for 6-aminoquinolyl-N-hydroxysuccinimidyl carbamate (AQC) synthesis were obtained from Sigma-Aldrich (Bangalore, India). Acetone was obtained from Fisher Scientific. Solid phase extraction cartridges (Strata-X, 8B-S100-TAK) were obtained from Phenomenex, Inc. (Hyderabad, India). High purity MS grade solvents (methanol, acetonitrile, and water) were obtained from Merck Millipore (Merck Millipore India Pvt. Ltd., Bangalore). Deuterated internal standards: L-serine-2,3,3-d3, L-glutamic-2,3,3,4,4-d5 acid, L aspartic-2,3,3-d3 acid, L-histidine-d3 (α-d1; imidazole-2,5-d2) HCl, L-tryptophan-2′,4′,5′,6′,7′-d5, L-4-hydroxyphenyl-d4-alanine-2,3,3-d3 (tyrosine), 4-aminobutyric-2,2,3,3,4,4-d6 acid (GABA-d6), histamine-α,α,β,β-d4, serotonin-α,α,β,β-d4, L-dopa-2,5,6-d3, 2-(3,4-dihydroxyphenyl)ethyl-1,1,2,2-d4-amine-HCl (dopamine-d4) and tryptamine-α,α,β,β-d4 were obtained from CDN Isotopes (Quebec, Canada). The deuterated internal standards 2-(4-Hydroxyphenyl)ethyl-1,1,2,2-d4-amine HCl and beta-Hydroxytyramine (α-d2, β-d1) HCl were supplied by Medical Isotopes, Inc. (USA). The purity of all analytes and deuterated internal standards was ≥98%. Glass beads and the bead beater were purchased from BioSpec

### 4.2 Animals and feeder training

Honey bee colonies (*Apis mellifera*, N = 6) were located inside the NCBS campus, Bangalore, India. Foragers were trained from the hive to an unscented sugar water feeder (concentration 1.75 M). The feeder distance was gradually increased to 300m from the hive over two days and foragers were trained along a road surrounded by dense vegetation in the neighboring UAS-GKVK Campus, Bangalore, India (Chatterjee et. al., 2019).

### 4.3 Behavior Experiments

Two kinds of behavior experiments were performed. In the first behavior experiment (BE 1), foragers (n = 10-12) were color-marked individually prior to the day of the experiment (Chatterjee et. al., 2019). On the day of the experiment, marked foragers were allowed to visit the feeder at 300m for 1.5 h (foraging phase). The feeder was then removed and kept hidden away from the reach of the foragers for another 1.5 h (search phase; Figure 1A). In the second behavior experiment (BE 2), the feeder was presented initially for 1.5 h (foraging phase) for marked foragers (n = 50-60) to visit. The feeder was then removed for 1 h (search phase) and reinstalled at 300m for another 1.5 h (revisit phase).

Colonies used for BE 1 (N = 2, 2015 and 2020) were housed in a glass observation hive located within a wooden hut devoid of any external illumination (Chatterjee et al., 2019). In case of BE 2 (N = 1, 2019) the colony was housed in a Langstroth hive box and placed under the shade of a tree next to the site of the hut. Experiments were performed in the summers between the months of May – July each year.

### 4.4 Collection Experiments

Three collection experiments (CE 1-3) were performed. During each experiment, color-marked foragers (n = 50-60) initially visited the feeder (concentration 2 M) for 1.5 h (foraging phase). At the end of the foraging phase (last 10 min), color-marked outbound foragers from the hive opening were captured in plastic tubes and flash-frozen in liquid nitrogen. The feeder was then removed and kept hidden during the search phase (1-1.5 h). Foragers flying out did not find the feeder at 300m and came back to the hive. Marked individuals were caught while flying out after 10-90 mins of feeder removal. The feeder was reintroduced at 300m (revisit phase) and foragers readily continued foraging at the feeder for another hour. Revisiting individuals were captured at the hive entrance making outbound flights, provided each made 3-4 trips to the feeder (Figure 2A).

Colony used for CE 1 (N = 1, 2017) was housed in a glass observation hive located within a wooden hut devoid of any external illumination (Chatterjee et. al., 2019). In case of CE 2-3 (N = 2, 2018 and 2019), the colonies were housed in a Langstroth hive box and placed under the shade of a tree next to the site of the hut. Experiments were performed in the summers between the months of May - July each year. For details about numbers of bees analyzed, see supplementary Table S2.

### 4.5 Monitoring search flights

For BE 1 and CE 1-3, a video camera (Sony HDR-CX220V Tokyo, Japan) was mounted at the hive entrance. Experimenters recorded the number and duration of search flights made by individuals from the time of exit and entry of the foragers in and out of the hive from video playback. Video recordings were done at the feeder for BE 2 during foraging and revisit phase to monitor the rate of bees arriving at the feeder.

### 4.6 Brain dissections

The foragers collected in liquid nitrogen were transferred to a −80 ° freezer. Individual bee brains were dissected on dry ice into two different parts, the optic lobe pair (OL) and the region of the brain containing the mushroom bodies and the central brain (CB) (Figure S3). The brains were dissected out within 6 min and were never allowed to thaw. The trachea covering the brain becomes a thin film that can easily be brushed off without damaging the brain. We did not rinse the brain in any liquid, in order to preserve the tissue integrity and prevent the degradation of biogenic amines. Brains were dissected and prepared for subsequent mass spectrometric analysis prior to the completion of video analysis and classification of individual bees based on their behavior. Samples were excluded from statistical analysis only if during the video analysis, the bee could not be identified because the marking was blurred, or because of contrast and brightness issues or the bee was upside down, etc.

### 4.7 Selection criteria for sample processing

From the number of bees that were collected for a given experiment, we made an effort to include equal numbers of samples from all behavioral groups. The selection of samples for MS processing and analysis was done based on the time of capture during the collection experiment (foraging, searching and revisiting) whereas the final classification of the behavioral group was done only after time-consuming video analysis. Statistical analysis was done on F, S and R as well as F, FS, S1, S2 and R depending on the behavioral phenotype identified in the video analysis. Samples were excluded from statistical analysis if the bee could not be identified during the video analysis because the marking was blurred, or because of contrast and brightness issues or the bee was upside down etc. As a consequence of the delayed behavioral analysis, several of the samples for which we had mass spectrometric data could not be incorporated in the final statistical analysis.

### 4.8 Mass spectrometry of neurotransmitters

For mass spectrometric measurements, brain samples were prepared as in Ramesh and Brockmann, 2019. Briefly, to the vial containing the individual bee brain part, 100 μL of 0.5 mm glass beads, 190 μL of acetone containing 0.1% formic acid, 10 μL of 10 mM freshly prepared ascorbic acid, and 10 μL of 0.5 μg/mL internal standard mixture was added. Five microliters of a 0.5 μg/mL serotonin and tryptamine mixture was added to each sample (spiked) to aid quantification of these low ionizing and high matrix suppressed compounds. A bead beater was used to homogenize the samples, and the supernatant was collected in a new vial and lyophilized. For the derivatization procedure, 80 μL of 200 mM borate buffer and 10 μL of 10 mM ascorbic acid was added to the lyophilized extract, mixed well and 10 μL of a freshly prepared solution of 10 mg/mL AQC was added. The reaction was incubated at 55 °C for 10 min and stopped by the addition of 3 μL of 100% formic acid.

MS grade water (500 μL) was then added to the samples, and samples were loaded onto activated and equilibrated RP-SPE columns and washed twice with 1 mL of water containing 0.1% formic acid. Elution was done with 1 mL of ACN-MeOH (4:1) containing 0.1% formic acid and lyophilized and stored at −20 °C until injection into the instrument. Samples were reconstituted in 50 μL of 2% ACN containing 0.5% formic acid. The calibration curves range for each compound were made according to the abundance in the biological matrix and were the same as in Ramesh and Brockmann, 2019 and are given in the supplementary data file S1. Comparison of instrument responses for calibration curves over the three years of measurements are given in Figure S7.

A Thermo Scientific TSQVantage triple stage quadrupole mass spectrometer (Thermo Fisher Scientific, San Jose, CA, USA), connected to an Agilent 1290 infinity series UHPLC system (Agilent Technologies India Pvt. Ltd., India) was used for the neurochemical quantification. The column oven was set at 40°C, and the autosampler tray at 4°C. The mobile phase solvent A was 10 mM ammonium acetate containing 0.1% formic acid, and solvent B was ACN containing 0.1% formic acid. A C-18 column (2.1 mm × 100 mm, 1.8 μm, Agilent RRHD ZORBAX) fitted with a guard column (2.1 mm × 5 mm, 1.8 μm Agilent ZORBAX SB-C18) was used. The LC gradient was as follows: (2% B at 0 min, 2% B at 3 min, 20% B at 20 min, 35% B at 25 min, 80% B at 25–27 min, 2% B at 27–35 min) at a flow rate of 0.2 mL/min. The MS operating conditions were as follows: 3700 V spray voltage (positive ion mode); 270 °C capillary temperature; 20 (arbitrary units) sheath gas pressure; 10 (arbitrary units) auxiliary gas; argon collision gas. The S lens voltage and collision energy were as given in Ramesh and Brockmann, 2019, and are also provided in the supplementary data file S1. Quantification was done using the Xcalibur software version 2.2. Mass spectrometric measurements of the brain samples were done at the NCBS in-house facility.

### 4.9 Sample stability and storage

Frozen and dissected brains can be stored in the −80°C deep freezer for up to a year and processed and freeze-dried samples can be stored in the −20 freezer for up to 2 weeks. Known amounts of deuterated internal standards are added before any kind of sample processing is done, to normalize for any sample loss throughout. Compounds remain highly stable over multiple weeks under lyophilized conditions, post derivatization. In aqueous solutions, the AQC derivatized products start degrading, but the samples were reconstituted only just before injection. Under aqueous conditions, 80% of the ion intensity is still present after 48 hours. Validation of this method has also been done previously by others (Natarajan et al., 2015).

### 4.10 Statistical Analysis

#### 4.10.1 Classification of outbound trips

Trips of the foragers were classified based on the foragers’ knowledge whether the feeder was present or not into F, FS and S (Table S1). In both F and FS, a forager flying out of the hive had the information that the feeder was present whereas in S(s) the forager had the information that the feeder is no more present.

#### 4.10.2 Changes in hive-to-hive duration and hive stays

For behavior experiment BE 1, we wanted to know if the removal of the feeder led to changes in (a) hive-to-hive duration for an outbound flight and (b) duration of the hive stays between two outbound trips. We used a generalized linear mixed-effects model (GLMM) with a Gamma error distribution, considering the individual identity of the bee as a random factor (individual effect). We compared (a) hive-to-hive duration (F, FS and S1-4) and (b) duration of hive stay following outbound flights (HF, HFS, HS1-3) using a generalized linear hypothesis test (GLHT). P-values were corrected for multiple testing using single step adjustment. Distribution structures of the data were determined prior to model building by comparing AIC values as a goodness of fit criteria. Models were used separately for the two colonies.

Determination of distribution structures, GLMM and GLHT were done using the “fitdistrplus”, “lme4” and “multcomp” packages respectively in R version 4.0.2.

#### 4.10.3 Search behavior sequence and cluster analysis

A combination of sequence and cluster analysis (Lowe et.al., 2020) was used to identify common search behavior patterns among individual foragers following feeder removal in the behavior experiment BE 1. First, consecutive search flights and in-between hive stays for a forager were arranged as a search behavior profile. Each profile started with the exit of the bee for FS (t= 0 min) and was terminated at 120 min. When a bee came back from a search flight and did not appear outside until the end of the observation period, the time the forager spent inside the hive was counted as her final hive stay. For example, if a bee made 4 search flights, the search behavior profile would include the search behavior states: “FS - HFS - S1 - HS1 - S2 - HS2 - S3 - HS3 - S4 - HS4” where HS4 is the final hive stay (i.e., final state in the sequence). Next, the duration a forager spent in a given state was rounded to its nearest whole number (minimum duration of stay in a given state is 1 min). Then, each search behavior profile (SPELL format) was converted into a search behavior sequence (STS format). The whole observation period of 120 min was divided into bins of 1 min and a forager occupied a given state for 1 min and moved into a new one or continued to be in the same state in the next 1 min depending on the search behavior profile. The sequences of search behavior states occupied by foragers were stored chronologically in a matrix (rows for every forager and the columns for a given state, see also supplementary data file S1 for details).

Finally, we calculated a distance matrix, i.e., the distances between all pairs of search behavior sequences using the optimal matching distance metric. The metric used an insertion/deletion cost of 1 and substitution cost using transition rates (min = 0 for identical substitution and max = 2 for a transition not observed) between observed states in the search behavior sequence. Individuals with common search behavior sequences were grouped by hierarchical agglomerative clustering using Ward’s D2 clustering criterion based on the distance matrix computed earlier. The optimal number of clusters was determined by selecting the maximum average silhouette width (Figure S2A; Levshina, 2015).

For collection experiments (CE 1-3), the search behavior profile for an individual making 2 search flights before being captured included the search behavior states: “FS - HFS - S1 - HS1 - S2 - HS2” and HS2 was the final state in the sequence. A combination of sequence and cluster analysis was further done similar to behavioral experiments as mentioned before to identify common search behavior patterns among individual foragers following feeder removal (see also additional data file S1 for details). Foragers in CE 1-3 (n=41) were grouped into three clusters (I-III; Figure 4A) based on the similarity in duration of stay in given states in their search behavior profile (Figure 4B; see also Figure S2B for optimum number of clusters). Cluster I consisted of 7 FS bees (n=6, 2018 and n=1, 2019) which had their final stay in the hive (HFS) more than 20 mins. (Figures 4A and 4B). Cluster II had 2 S1 bees (n=1, 2018 and n=1, 2019) which stayed in the hive (HS1) for more than 35 mins before being captured. The biggest group, Cluster III consisted of S1, S2 and S3 bees (n=15, 2017, n=15, 2018 and n=2, 2019) which all had their final stay in the hive (HS1, HS2 and HS3 respectively) for less than 20 mins.

Cluster III was further subdivided into 5 subgroups post-hoc (IIIa-IIIe; Figures 4A and 4B). Subcluster IIIa comprises 13 FS bees which had their final stay in the hive (HFS) no longer than 14 mins. Subcluster IIIc (n=4) had all S2 bees but one individual (BeeID: C36) forming an outlier in subcluster IIIb with longer FS and search flights (S1 and S2 more than 18 mins). Subclusters IIId (n=2) housed S1 bees which stayed in the hive (HS1) longer than 15 mins (but less than 35 mins) before being captured. Finally, subcluster IIIe (n=12) housed the rest of the S1 bees which spent time in the hive (HS1) less than 9 mins before they were captured (see also supplementary data file S1 for details). This clustering further led to comparing neurotransmitters among individuals.

The sequence analysis was performed using the package “TraMineR” and agglomerative clustering was done by using the “agnes” function from package “cluster” in R version 4.0.2.

To help with the neurochemical data analysis, the FS and S bees without full behavioral data (7 bees) were manually classified into the identified clusters. For this purpose, the number of search flights and the total amount of time a bee experienced a loss of feeder before being caught were used in addition to incomplete flight duration and hive stay data. The details of the clustering criteria are given in the supplementary data file S1.

#### 4.10.4 Quantification of foragers dynamics at the feeder

For BE 2, the total number of marked foragers at the feeder was counted every 2 minutes (Figure S1). We asked if the rate of foraging was different during the foraging and revisiting phases. We fit non-linear growth curves to the number of bees at the feeder every 2 min to evaluate and compare the rate of foraging:

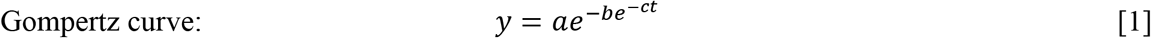

where a = carrying capacity (maximum number of bees), b = sets the displacement along the time-axis, c = growth rate, t = time in minutes and the inflection point (I) is given as equation 2 (Jukić et. al., 2004).

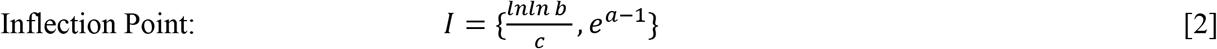

Gompertz curve fitting was done using the “nlsfit” function from package “easynls” in R version 4.0.2.

#### 4.10.5 Analysis of the mass spectrometry results

Neurochemical analysis was done using linear mixed effects models from the “lme4” package. Satterthwaite’s t-tests from the “lmerTest” package were used to estimate significance values from the models. Analysis was done with the amount of neurochemical as the response variable, and the appropriate behavioral response/behavioral group as the fixed variable. The MS batch was used as the random effect. The formula (given in the R syntax) used for the model is as follows:

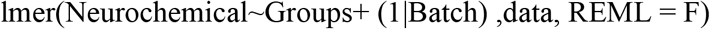

Post-hoc tests were done using the emmeans function from the “emmeans” package using the following code:

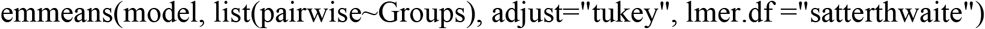

Plots were drawn using the “ggplot2” package. For some plots, for visual purposes, the neurochemical values were scaled with respect to the MS batch and the experimental repeat. PCA analysis was done using the “prcomps” function in the “stats” package of R. The package “ggbiplot” was used, with a minor adjustment for visual purposes, to plot the prcomp results.

## Supporting information

Supplementary Datafile

## 6 Author Contributions

Conceptualization A.C., D.R. and A.B.; Field experiments and observations, A.C. and D.B.; Brain Dissections, D.B.; Behavior analysis, A.C.; Mass spectrometric measurements and analysis, D. R.; Writing, A.C. and D.R and A.B.; Supervision, A.B.

## 8 Acknowledgments

The authors would like to thank student interns A. Sengupta, A. Suryanarayanan, S. Chakraborty, D. Chowdhury and A. Chakrabarty for helping with the behavior experiments. The authors also thank the NCBS Mass Spectrometry facility.

## 9 Supplementary Material

Supplementary Material is available with this manuscript.

## 1 Data Availability Statement

The original contributions presented in the study are included in the article/supplementary material, further inquiries can be directed to the corresponding author/s.

