## Supplementary Figures and Tables for "Search behavior of individual foragers involves neurotransmitter systems characteristic for social scouting"

### *Supplementary Material*

#### 1 Supplementary Figures and Tables

##### 1.1 Supplementary Figures

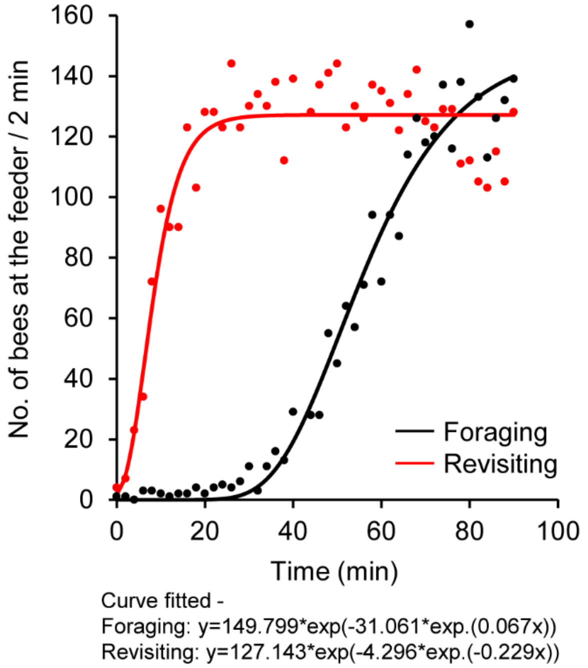


**Supplementary Figure 1.** Dynamics of foragers visiting the food source before removal (black) and after re-establishing the feeder (red). Foragers searching for the missing feeder started visiting at a faster rate when the feeder was re-established compared to when the feeder was opened for the first time on that day.

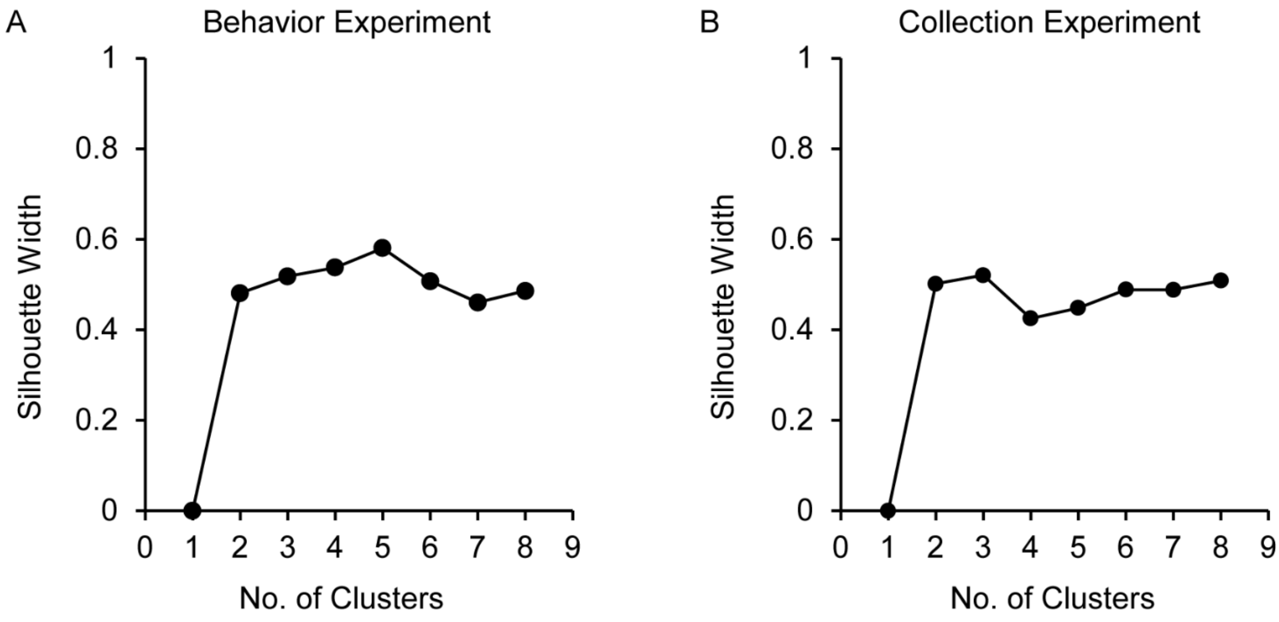


**Supplementary Figure 2.** Silhouette width for determining the number of clusters. (A) Behavior Experiment, (B) Collection Experiment. The maximum silhouette width determined the number of clusters.

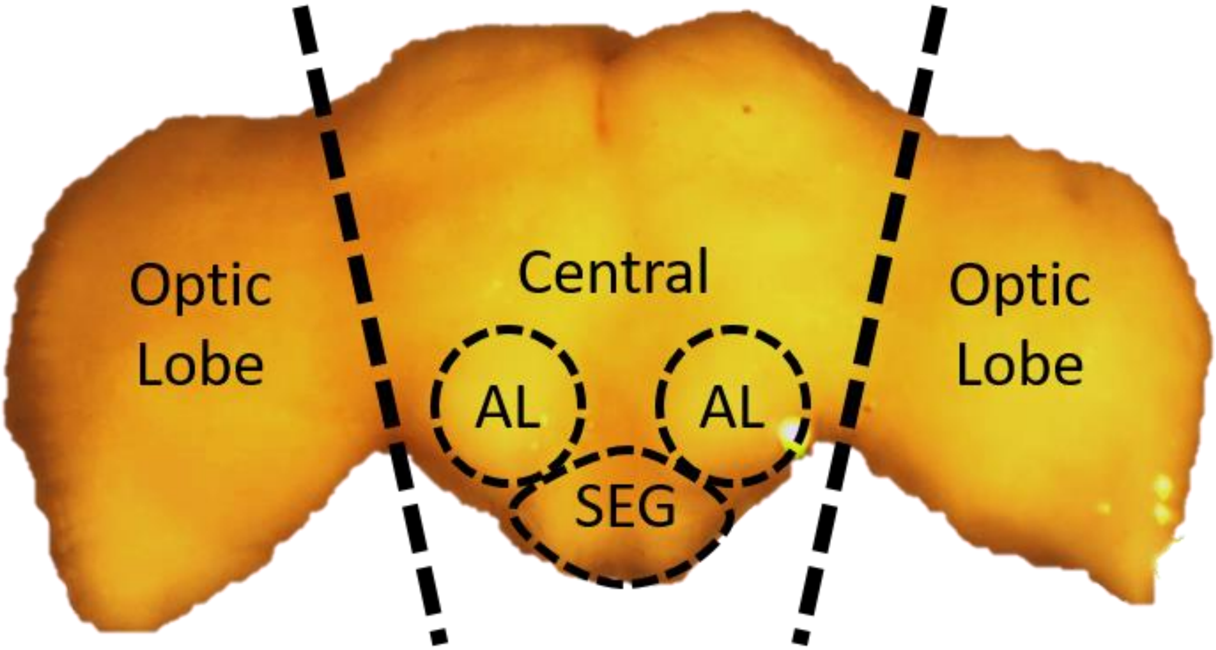


**Supplementary Figure 3.** Honey bee worker brain (frozen) showing regions and breakpoints used in this study for dissections.

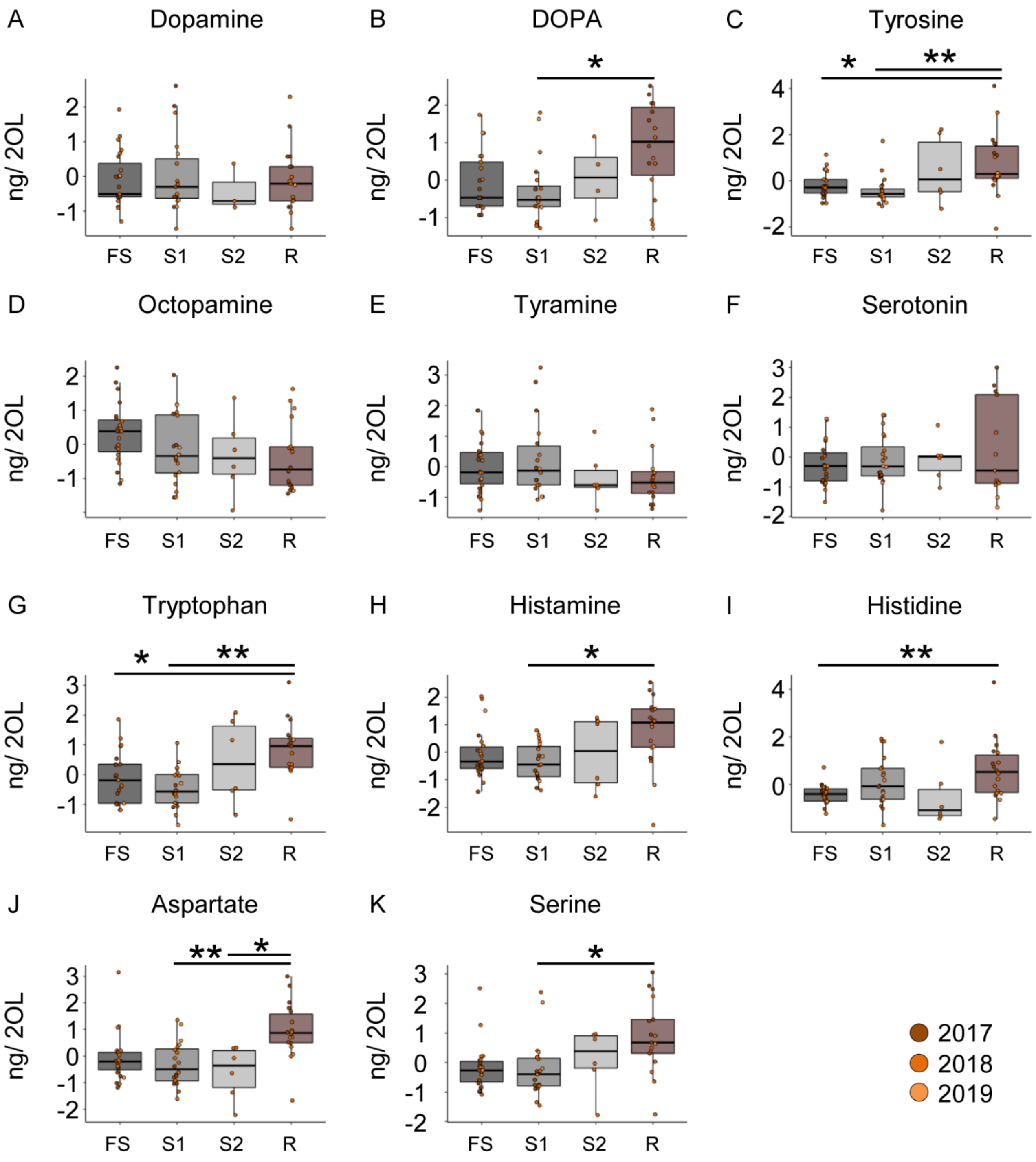


**Supplementary Figure 4.** Foragers that initiated foraging have the highest neurotransmitter titers in the OL. The abrupt increase in transmitter content seen in the OL of foragers that initiated foraging are independent of the number of search flights performed by the bees.

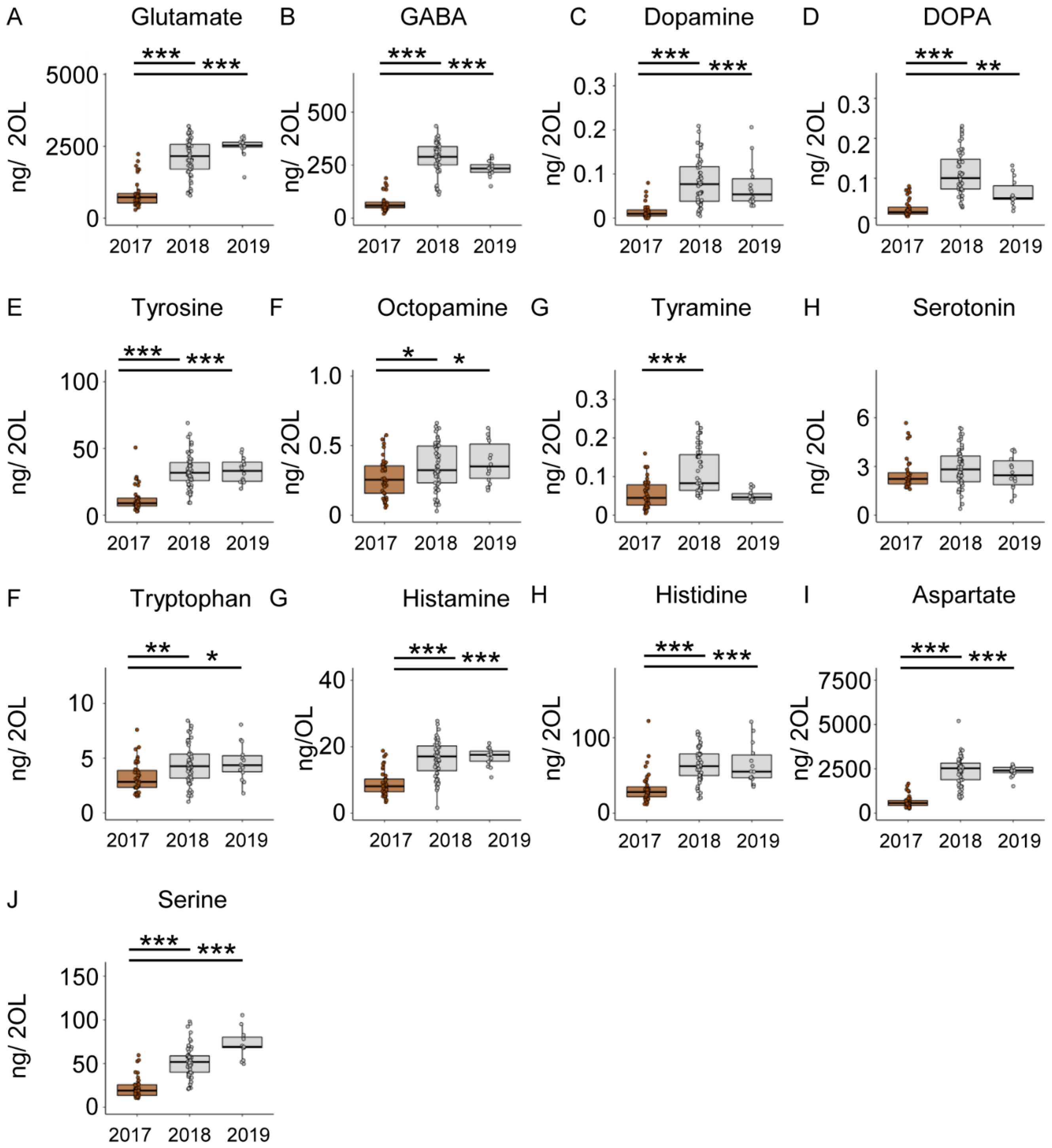


**Supplementary Figure 5.** Colony dependent variation in neurotransmitter titers in OL. Except for serotonin (H), all other transmitters showed significant differences due to the colony identity. In addition, foragers from the CE 1 contained significantly lower amounts of all changed transmitters. Only the differences between 2017 and the other two years are shown.

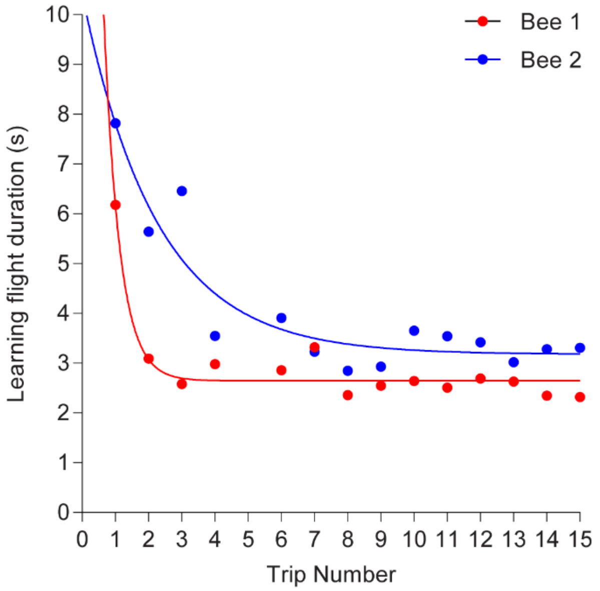


**Supplementary Figure 6.** Learning flights. A learning flight is a specific flight pattern the forager performs after leaving the feeder prior to the vector flight to the hive. It is described as a “turn back and look” event during which the forager departing from the feeder makes consecutive circles in the air with increasing radius and height for ~ 8-10 sec to examine the location to acquire the visual and landmark information to guide her return (Lehrer, 1991; Wei et. al., 2002; Wei and Dyer, 2009). This behavior is an example of “backwards conditioning” where learning is followed after the reward is received and is essential since bees could not find their way back to the hive without a learning flight (Lehrer, 1993). In a single shift experiment (200m-300m), the durations of the learning flights were analyzed for two foragers (red and blue). Learning flight duration decreased with successive trips to the new feeder location (unpublished data). We also found foragers making learning flights when they departed from a re-established feeder at the same location (personal observation) which might indicate re-learning the feeder location with respect to the environment around it.

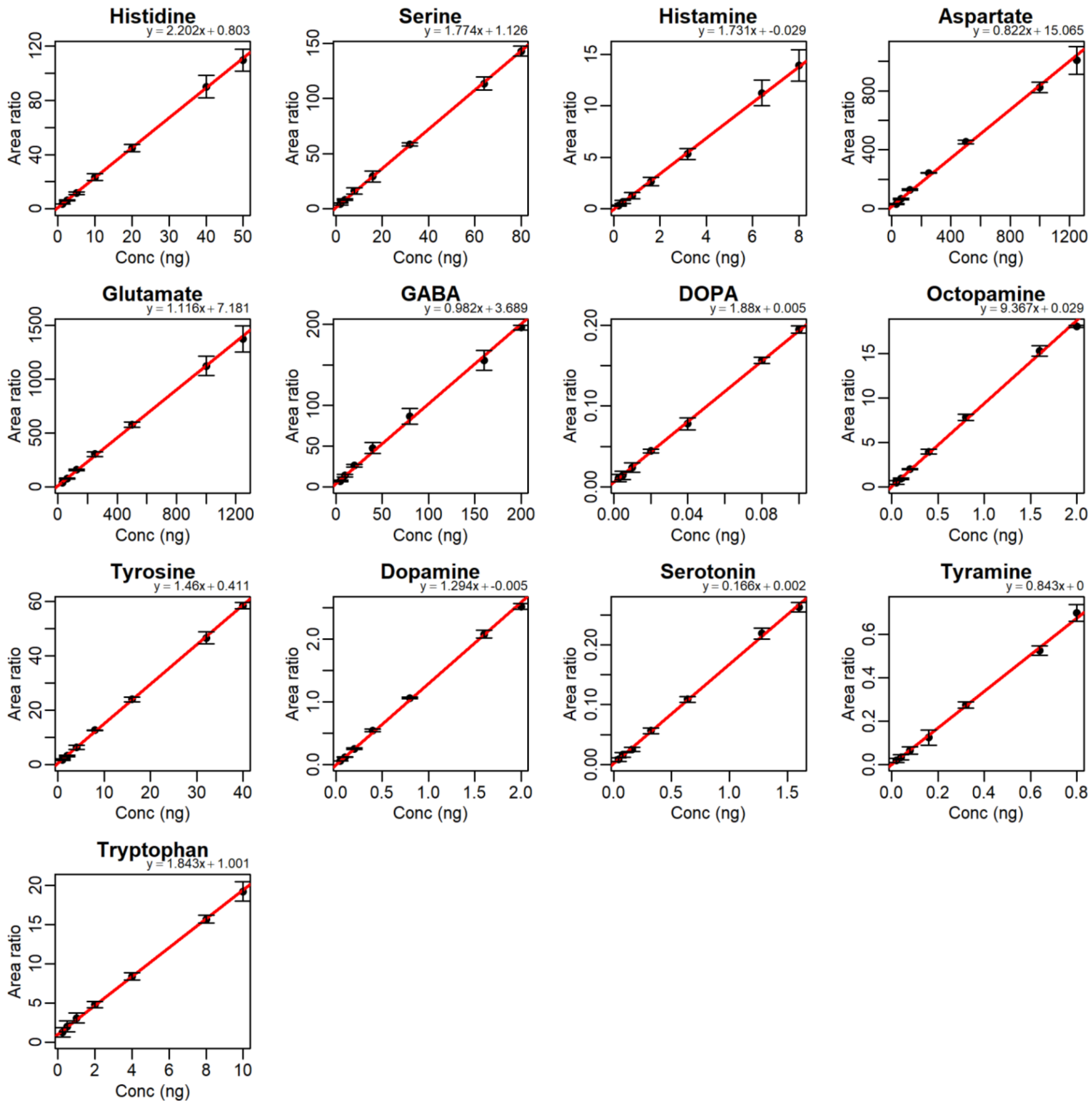


**Supplementary Figure 7.** Calibration curves.

#### 1.2 Supplementary tables

**Table S1:** Classification of outbound and inbound flights

| **Classification** | **What do the foragers know?** | |
|---|---|---|
| | **Outbound flight** | **Inbound flight** |
| Foraging (F) | Present | Present |
| Foraging + Search (FS) | Present | Absent |
| Search Flight (SF) | Absent | Absent |

**Table S2:** Summary of collection experiments

| Colony | Year (Experiment) | Number of bees analyzed (CB/OL) | | | |
|---|---|---|---|---|---|
| | | F | FS | S1-S2 | R |
| 1 | 2017 (CE 1) | 11/12 | 11/12 | 6/7 | 8/8 |
| 2 | 2018 (CE 2) | 23/23 | 11/10 | 15/16 | 8/8 |
| 3 | 2019 (CE 3) | 7/8 | 1/1 | 3/3 | 2/2 |

**Table S3:** Comparative time scale associated with search behavior

| Comparative time scale associated with search behavior | Reference |
|---|---|
| Hive to hive trip to shut feeder (60 m*): < 5 min<br>Dance guided search trip: 17 ± 11 min (mean ± sd) | Seeley, 1983 |
| Homing flight duration of foragers captured at a feeder (10 m) and displaced to a release site (200 - 250 m): 61.78 ± 84.36 min (mean ± sd), range: 2.13 – 121.43 min | Reynolds et. al., 2007a |
| Search flight duration of foragers for a missing feeder (210 m): 4.49 ± 2.44 min (mean ± sd) | Reynolds et. al., 2007b |
| First return trip of foragers to the hive following downshift of reward (within a flight cage 35 m long): < 25 min<br>Time spent within hive: 25.6 ± 2.8 (mean ± se) | Townsend-Mehler and Dyer, 2012 |
| Visit-persistence to an emptied feeder (20m, 450m): 4.29 ± 4.47 trips (mean ± sd), range: 0 – 25 over a 6 hour /day | Al Toufailia et. al., 2013 |
| Hive to hive duration for a missing feeder (300 m): Foraging + Combined foraging/search flight (FS): 9.2 ± 4.5 min<br>Search flights (S): 8.31 ± 3.5 min (mean ± sd) | Chatterjee et. al., 2019 |

*The distance of the feeder is in meters from the hive.
